# Targeted Sequencing of Alzheimer’s Disease Genes in Gujarat Cohort

**DOI:** 10.1101/2022.04.14.488366

**Authors:** Urvi Budhbhatti, Chaitanya Joshi, Ajay Chauhan, Deeptiben Bhatt, Chirag Parmar, Vishalbhai Damani, Amrutlal Patel

**Author notes:** Corresponding authors: (AP).

## Abstract

Alzheimer’s Disease (AD) is the most common type of dementia affecting cognitive and behavioural functions. It’s a complex disease that results from the modest effect of gene interaction and environmental factors, as a result of which its exact pathogenesis is still unknown. In the present study, we have performed an association study of exonic regions of 94 genes with AD using the Haloplex target enrichment method and genotyping of 4 genes NOTCH4(rs367398), NEP(rs61760379), APOE (rs121918398), and ATP7B(rs1801243) using PCR RFLP technique. In this study, we screened 32 AD cases and 11 control samples. We have identified 19 variants assigned to 16 genes that were significantly associated with AD with a p-value < 0.05. However, no variants have passed multiple test corrections except NOTCH4/rs367398 genotyped using the PCR-RFLP approach. The functional enrichment analysis of the significant genes for the disease indicates that the genes are significantly involved in AD. Also, functional enrichment analysis for biological processes using ClueGo and network analysis using the STRING database indicated that the genes are directly and indirectly playing a role in AD pathogenesis. The eQTL analysis of the variants with GTEx dataset indicate that out of 19 variants ACHE/rs1799806 and NOTCH4/rs367398 variants is showing significant eQTL for the neighbouring genes for different brain regions and whole blood. Regardless, additional studies with a large cohort are required to establish the role and prevalence of these variants in the local population of Gujarat.

## 1. Introduction

Alzheimer’s disease (AD) and other forms of dementia presently accounts for 40-50 million cases worldwide[1]. Dementia affects 5.3 million individuals in India, and the number is expected to rise substantially as living and health standards improve, leading to a longer life expectancy. By 2035, it is expected that the number of dementia cases is likely to be doubled[2]. Clinically, AD is a progressive neurodegenerative disorder that leads to severe cognitive impairment subsequently requires full-time medical care. Apart from the significant effect of AD on patients and primary caregivers, the high expenditures involved with dementia care and treatment place a significant burden on society and public health[3].

The genes amyloid precursor protein (APP), presenilin1 (PSEN1), and presenilin2 (PSEN2) have been linked to Early-onset autosomal dominant Alzheimer disease (EOAD). However, atypical highly penetrant mutations in APP, PSEN1, and PSEN2 account for only a small percentage of EOAD (5-10%), leaving a large number of patients with inexplicable EOAD genetics, inferring that other susceptibility genes were yet to be discovered[4]. Large-scale GWAS and subsequent meta-analysis found a collection of 20 candidate genes for Late-onset Alzheimer’s disease (LOAD) susceptibility, most of which are associated with APP metabolism, particularly with amyloid β-protein (Aβ) clearance[5]. Moreover, next-generation sequencing has revealed that the frequency of mutations in these genes varies dramatically among populations, with genetic background playing a profound impact in this variance[6]. Although various rationales incorporating genetic components have been proposed, precise aetiology is yet to be uncovered [7].

Considering the significance of mutational variations in specific pathways such as APP processing, oxidative stress and inflammation, and the dearth of data from the Indian population. The current study involves the basic association test using the targeted sequencing of 94 genes and analysis of 4 genes NOTCH4(rs367398), NEP(rs61760379), APOE (rs121918398), and ATP7B(rs1801243) using PCR RFLP approach that were implicated in AD.

## 2. Material and methods

### 2.1 Study subject

Samples of 32 AD cases were collected with their and their relative’s written consent from the Hospital of Mental Health of Gujarat (HMH-Ahmedabad, HMH-Bhuj, and HMH-Baroda). Samples were screened for 6 tests: neuropsychology test-Mini-Mental State Exam (MMSE), hemogram report, thyroid hormone test, Venereal Disease Research Laboratory (VDRL) Test, HIV I & II Test, Vitamin B-12 level, and Magnetic Resonance Imaging (MRI) report of the brain. The average age of the patient was 68.11 years, ranging from 52 to 79 years. All patients were selected with no family history of dementia. Meanwhile, an age-matched control group of 11 people was selected from the Gandhinagar and Ahmedabad areas nearby, consisting of 7 males and 4 females. Healthy volunteers with no clinical signs of neurological or psychiatric disorders and above the age of 60 years were considered for the control group. The present study was reviewed and approved by the Ethics Committee of HMH-Gujarat Institute of Mental Health (Ethics/Approval/GBRC/01-19).

### 2.2 Alzheimer’s disease panel

The Haloplex probes were designed using Agilent SureDesign software with the 94 genes that were screened after literature review, encompassing both previously recognized causal genes and those found in the preliminary review. These target regions include only exonic regions of the selected genes (Table S1). Probes were designed for the Illumina platform of 150 base pair chemistry. The design is anticipated to cover 99.79% of targeted genes.

### 2.3 DNA extraction

Qiagen’s QIAamp DNA blood mini-kit have been used to extract genomic DNA from peripheral blood collected in EDTA tubes as per the manufacturer’s procedure, and the quantity and quality were evaluated using Qubit DNA HS kit and 0.8% Agarose gel respectively.

### 2.4 Target enrichment method, Library Preparation, and NGS

Library prepared using the Haloplex HS target enrichment kit following the manufacturer’s protocol. The gDNA was digested with 8 pairs of restriction enzymes. Biotinylated Haloplex probes hybridized to the target region and direct its circularization. Circularized target fragments were then ligated and captured using Streptavidin beads. Captured target fragments were then PCR amplified and purified. The purified library was validated using the bioanalyzer and quantified using the Kapa library quantification kit. Mi-seq Illumina sequencer was used for the sequencing of the pooled samples.

Apart from targeted sequencing of 94 genes, we have also used PCR-RFLP approach to genotype 4 SNPs of NOTCH4(rs367398), NEP(rs61760379), APOE(rs121918398), and ATP7B(rs1801243) (Paper under communication).

### 2.5 Bioinformatics analysis

Initially, FASTQC was done to examine the quality of the reads and to determine the trimming parameters. Illumina adaptors and low-quality reads were trimmed using Trimmomatic v.0.39. FASTQC was done to ensure that adaptors and low-quality reads were removed from the trimmed Fastq file. BWA-MEM was used to align the trimmed paired forward and reverse Fastq data to the hg19 (GRCh37) human reference genome, generating SAM (Sequence Alignment Map) files. The SAM file is compressed into a BAM (Binary Alignment Map) file.

Picard tools were used to sort the BAM file and add read groups. Duplicates were not marked or removed, as was recommended for Haloplex[8]. The GATK (v4.2.0.0) BaseRecalibrator tool was used to perform Base Quality Score Recalibration (BQSR). The gVCFs of the individual samples were called using HaplotypeCaller ERC (Emit Reference Confidence BP_RESOLUTION) to call both variants and reference genotyping of the target regions. Individual gVCFs were then merged and joint genotyped using the CombineGVCFs and GenotypeGVCFs tools of GATK (v.4.2.2.0). The combined VCFs file was filtered for a minimum mean depth of 10 and was then converted to plink format for statistical association analysis. Coverage analysis for the target regions was done using bedtools.

### 2.6 Statistical analysis

Initially, Quality control (QC) was performed for the parameters such as minor allele frequency (maf), missing rate per individual (mind) and per SNPs (geno) were performed on the filtered dataset PLINK[9] (www.cog-genomics.org/plink/1.9/) with threshold values of 0.01, 0.2 and 0.2, respectively. The correlation between the SNPs and Alzheimer’s disease was evaluated in plink using the Fisher exact test with continuity correction. Plink was used to examine the statistical correlation of the PCR-RFLP genotyping data.

### 2.7 In-silico Analysis

Enrichment analysis using the Gene Annotation Tool WEB-based Gene SeT AnaLysis Toolkit (WebGestalt) for disease with the databases provided by DisGeNET. Non-random over-representation of genes from our gene list for Disease with False Discovery Rate (FDR) value < 0.05 was regarded as significant. ClueGO (v.2.5.8) has been used to infer the ontology and functional implications of 16 genes on AD metabolic pathways in this paper. ClueGO depict the non-redundant biological pathway with network genes cluster aggregated into a functional network using statistical analysis of existing Gene Ontology annotations.

Network analysis is widely used to study the functional association between genes. Functional gene-gene interactions analysis of 16 significant genes with the AD-associated top 10 genes (ACE, APP, ADAM10, GSK3B, HFE, APOE, MAPT, TREM2, PSEN1, and PLAU) selected from the DisGeNET database on the basis of the gene-disease association score was done using STRING database.

Expression Quantitative Trait Loci (eQTL) analysis is used to identify the susceptibility genes at the risk loci and the effect of SNPs in gene expression. SNP is linked to the particular gene if it is located within the gene or 1Mb of Transcription Start Site (TSS). Analysis of the significant SNP was recorded for the brain and blood tissue using GTEx v8 database. The genes with p value < 1E-03 was considered to be significant. Tissue specific expression pattern analysis of the genes that were significantly mapped to the pre-defined SNPs was done using GTEx database.

## 3. Result and Discussion

Using a targeted gene sequencing approach that was used to specifically enrich the target regions, we sequenced 43 DNA samples, comprising 32 patients with Alzheimer’s disease and 11 control subjects. We designed a sequencing panel comprising of 94 genes either directly or indirectly linked to AD.

On an average 99.76% reads mapped to the human genome. All genes achieved minimum coverage of 50X (Fig 1). The minimum coverage was observed for gene MS4A3 of 68.10X and maximum coverage was observed for the gene IRS1 of 235.99X.

**Figure 1:**
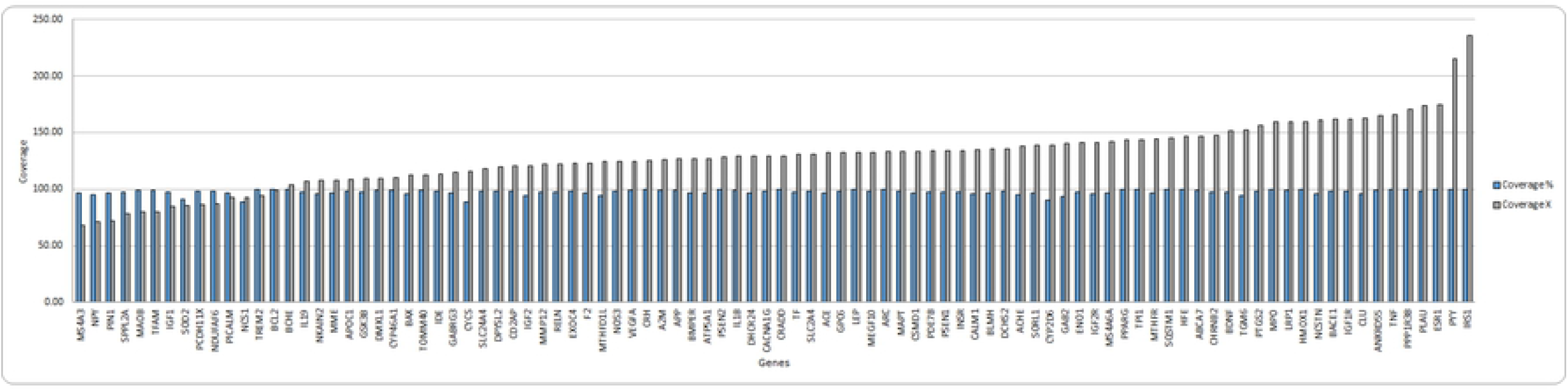
Average sequencing coverage achieved by 94 panel genes

Overall, 761 variants were detected, of which only 749 passed the minimum mean depth of 10 filter. Out of 749 variants, 38 were indels and 711 were SNVs. A quality check prior to statistical analysis filtered out 21 variants due to low maf threshold and 9 variants due to geno threshold. The average genotyping rate of the filtered dataset was 0.9972.

Table 1 shows the 18 variants annotated to 15 genes that were found to be significantly associated with AD with a p-value of 0.05. However, no variants passed multiple test corrections.

**Table 1:**
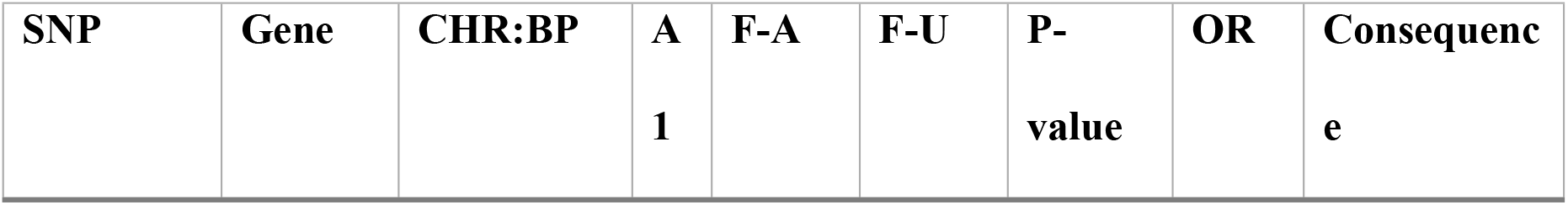
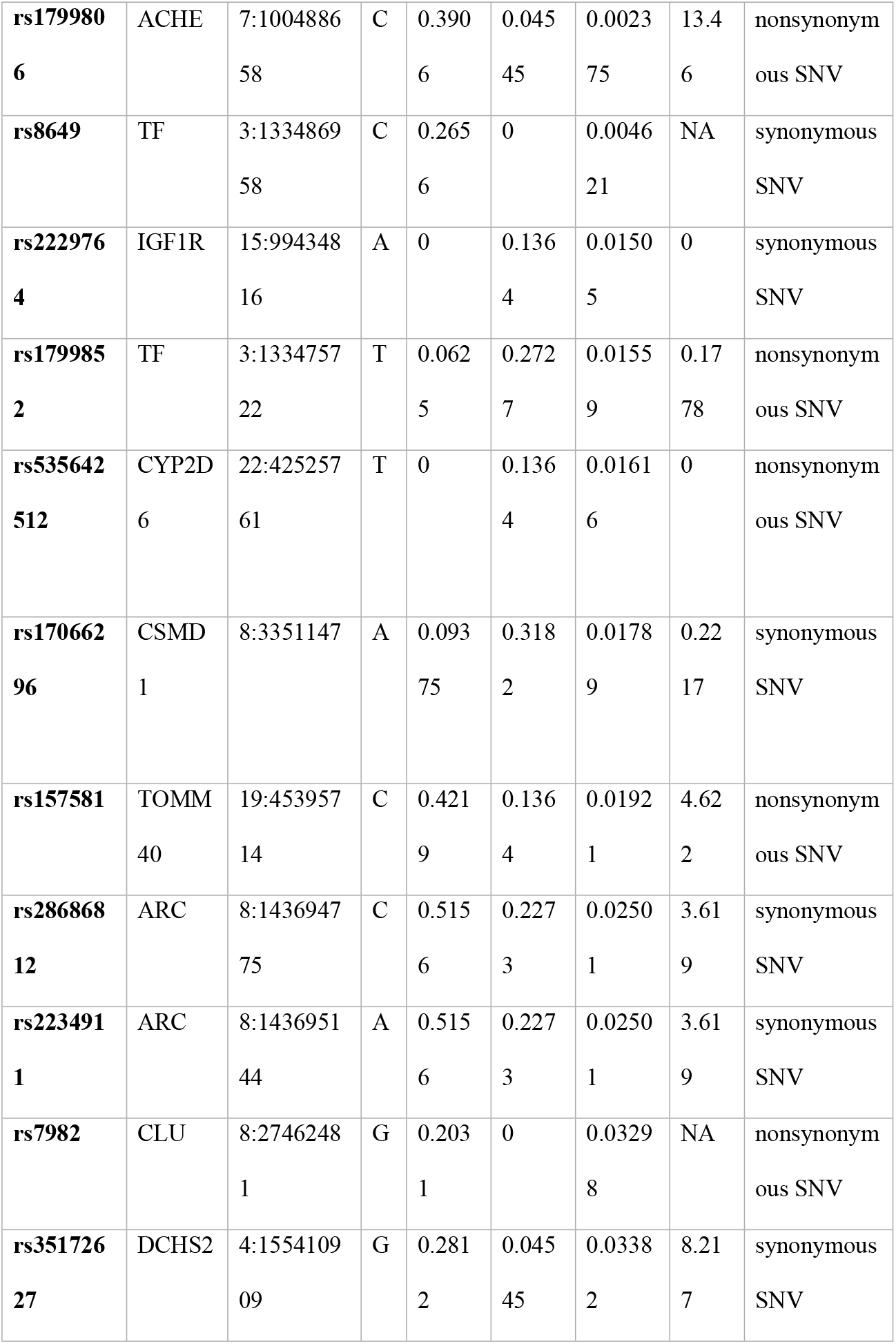
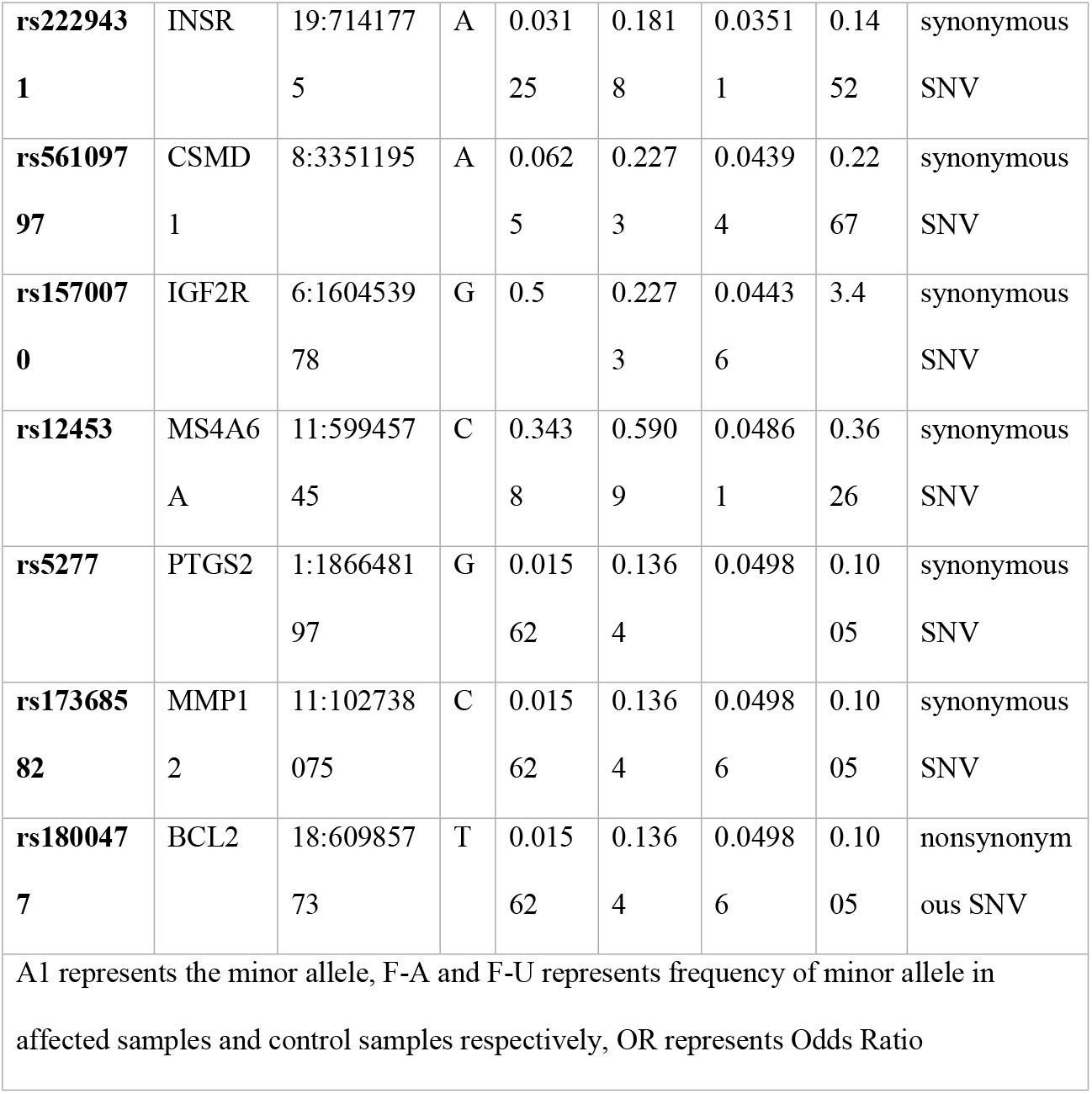
List of SNPs that were significantly associated with AD

Out of 18 variants, 12 were synonymous variants and 6 were non-synonymous. Although synonymous variants don’t alter the amino acid, studies have reported that synonymous variants may affect splicing, mRNA stability, and co-translational folding of protein[10]. It affects co-translational folding as the rare synonymous variants tends to translate more slowly than commonly used codon[11].

Statistical analysis of PCR-RFLP data shows that the NOTCH4/rs367398 a non-coding variant is significantly associated with AD (Table 2) with p value of 0.00576 and Odds Ratio(OR) of 5.1. In this cohort, however, no significant correlation of APOE/rs121918398, NEP/rs61760379, or ATP7B/rs1801243 was identified. NOTCH4/rs367398 variant is only variant significantly associated with AD that has passed multiple test corrections with bonferroni p value of 0.02305. In contrast to previous study conducted by Shibata N et.al., the NTOCH4/rs367398 is showing significant association with AD phenotype[12].

**Table 2:**
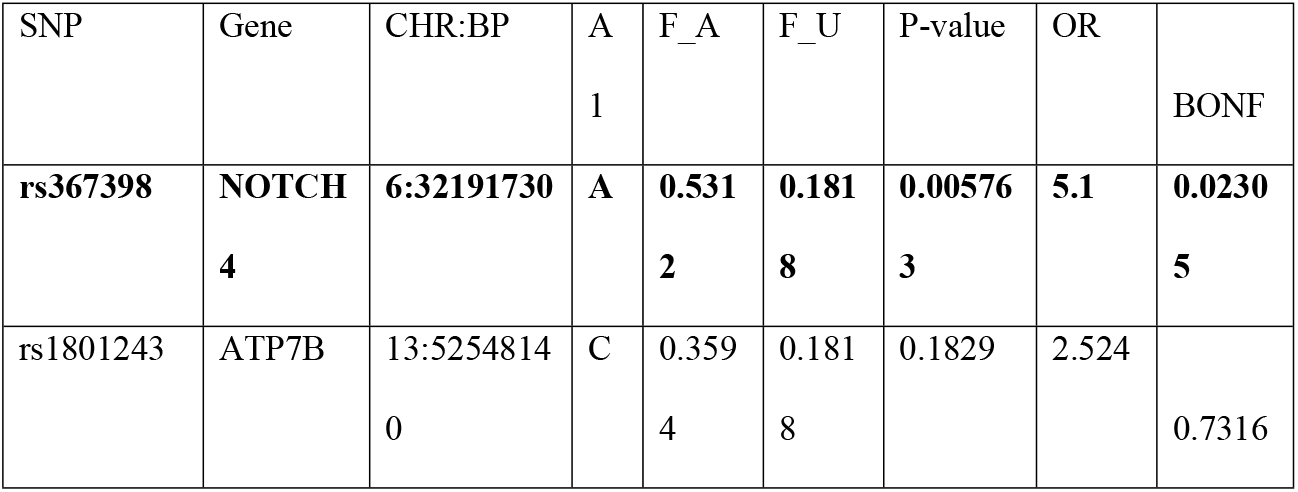
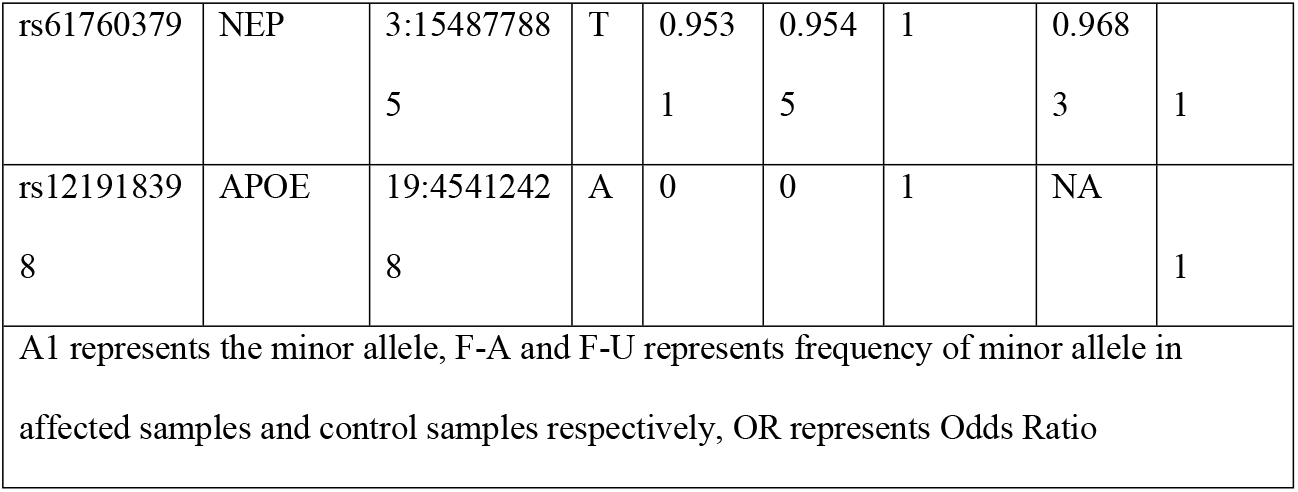
Results of Fisher analysis of PCR-RFLP genotype data using plink

According to meta-analysis of the two sequencing data by Fan KH et. al., the TOMM40/rs157581 variant significantly associated with AD (OR = 1.49; p = 4.04E-07)[13]. It has also been reported to have an impact on cognitive impairment[14]. The rs157581 variant is in linkage disequilibrium (LD) with the APOE gene variant rs405509 and has been shown to have a significant genome-wide correlation[15]. The CLU gene variant rs7982 has been linked to the baseline deposition of Aβ[16]. Contradict to the study conducted by Wang X et. al., the rs12453 variant of the MS4A6A gene has been shown to be associated with AD with a p value of 0.04861 and an OR of 0.3626[17].

### 3.1 In-silico analysis

Alzheimer’s disease (AD) is the most common form of dementia characterized by a progressive loss of episodic cognitive performance function, which leads to linguistic and visuospatial ability deficits, as well as behavioural abnormalities such as apathy, aggression, and depression[18]. The exact pathophysiology of AD is yet unknown because it results from the modest effect of gene interactions and environmental factors. The phenotypic effects of genes in an individual result from their interaction within gene regulatory networks and metabolic pathways.[19].

The enrichment analysis of the gene set for disease (Table 3) enriched the gene set for 8 diseases with an FDR < 0.05, with the highest enrichment of genes being 9 out of 15 for Alzheimer’s disease, with an FDR value of 8.97e-12. GO analysis helps to identify the functional gene-gene interaction within biological processes to evaluate the role of the genes and their implications in the AD pathway. The ClueGo analysis (Fig 2) revealed that the genes were majorly involved in nitric oxide metabolic processes, long chain fatty acid biosynthesis, synaptic transmission, apoptotic pathway in response to DNA damage, and insulin activated receptor response. Although the genes are not directly implicated in APP processing and Aβ clearance, the genes are indirectly correlated to Aβ processing. Nitric oxide metabolism is involved in neuro-toxic effects and reports suggest that nitrosative and oxidative stress are increased prior to Amyloid and Tau protein aggregation[20]. Oxidative stress plays an important role in AD and with age, susceptibility to lipid peroxidation increases, affecting neuronal homeostasis. Also, lipids are involved in cellular signalling and have an inflammatory role[21]. With an increase in age, DNA lesions increase and DNA damage and altered DNA repair pathways are proposed as risk factors for AD progression[22]. Also, insulin signalling affects Aβ clearance, synaptic function, and plasticity. As a result, changes in insulin signalling may increase the risk of Alzheimer’s disease[23].

**Table 3:** The list of diseases discovered to be significantly enriched in the enrichment analysis of the 16 significant genes

| Gene Set | Disease | P Value | FDR | Enriched Genes |
| --- | --- | --- | --- | --- |
| C000239<br>5 | Alzheimer's<br>Disease | 2.44E<br>-15 | 8.97e-12 | ACHE,TF,IGF1R,CYP2D6,ARC,CLU,INS<br>R, IGF2R,BCL2 |
| C002411<br>5 | Lung<br>diseases | 1.23E<br>-06 | 0.00225616<br>4 | IGF1R,INSR,IGF2R,PTGS2 |
| C004175<br>5 | Adverse<br>reaction to<br>drug | 2.54E<br>-06 | 0.00265719<br>7 | ACHE,CYP2D6,CLU,PTGS2 |
| C003056<br>7 | Parkinson<br>Disease | 2.89E<br>-06 | 0.00265719<br>7 | IGF1R,CYP2D6,INSR,IGF2R |
| C075234<br>7 | Lewy Body<br>Disease | 5.00E<br>-06 | 0.00367665<br>8 | IGF1R,INSR,IGF2R |
| C003634<br>1 | Schizophreni<br>a | 9.97E<br>-06 | 0.00610400<br>4 | ACHE,TF,CYP2D6,ARC,CLU,CSMD1,<br>PTGS2,BCL2,NOTCH4 |
| C023583<br>3 | Congenital<br>diaphragmati<br>c hernia | 3.12E<br>-05 | 0.01637424<br>5 | IGF1R,INSR,IGF2R |
| C145815<br>5 | Mammary<br>Neoplasms | 5.23E<br>-05 | 0.02400751<br>8 | ACHE,IGF1R,CYP2D6,PTGS2,BCL2,<br>NOTCH4 |

**Figure 2:**
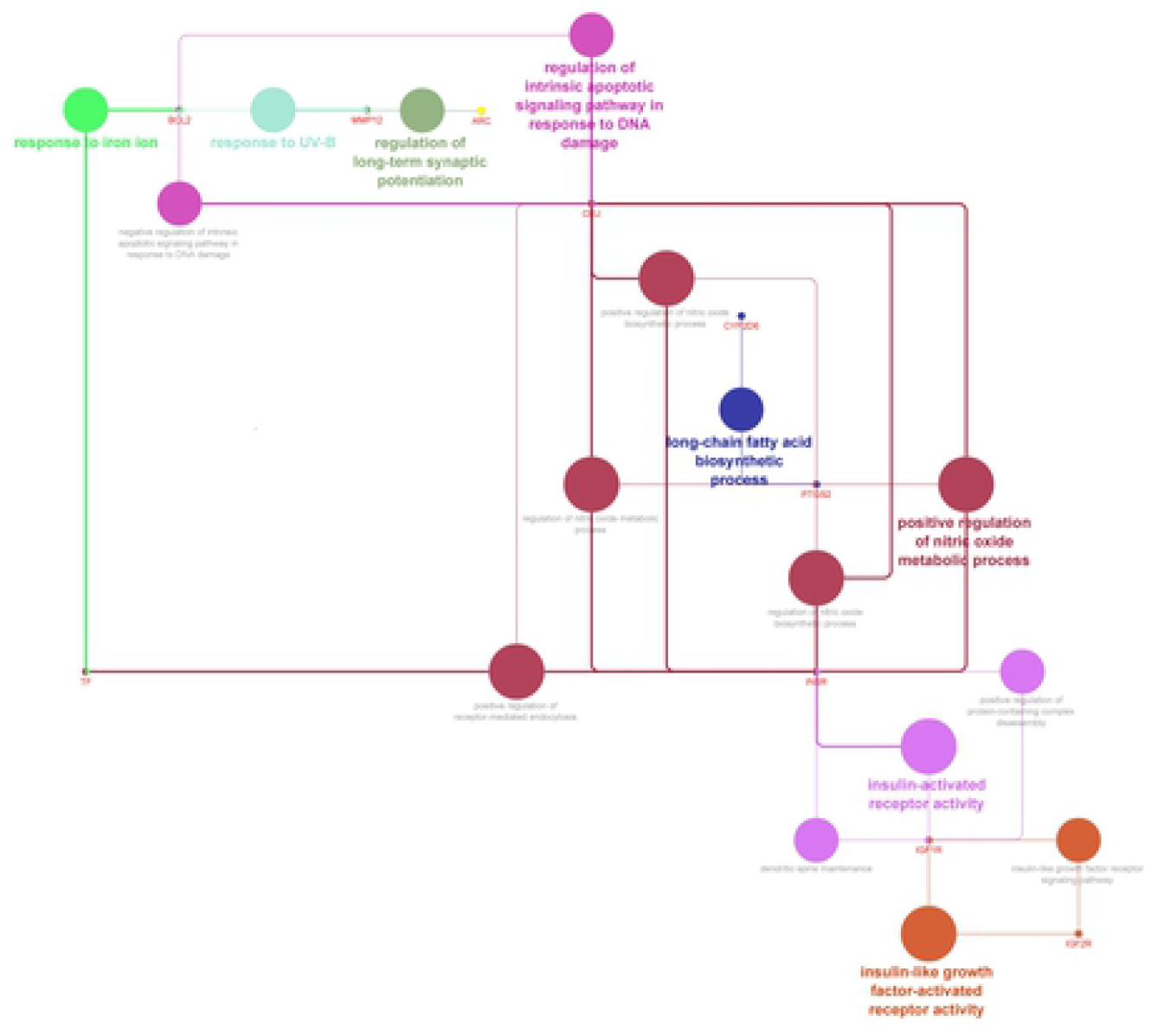
Enriched biological pathways that could play an indirect role in Amyloid metabolism and APP processing

Thousands of prevalent variants with high statistical correlations with phenotypic attributes have been uncovered through genome-wide association studies. Unfortunately, the functional implications of the majority of GWAS-implicated variants are mostly unexplored. The impact of genetic alterations on the genetic expression of one or more genes can be determined via eQTL analysis. In the present study we have done eQTL analysis using the GTEx v8 dataset[24]. The eQTL analysis revealed that out of 19 variants 7 variants is showing significant eQTL for different regions of brain and whole blood tissue (Table 4). Multiple neighboring genes have strong eQTL for ACHE/rs1799806. The SLC12A9 gene, which has a pvalue range of 3.00E-18 to 8.6E-0.6 in brain regions and a p-value of 7.00E-14 in whole blood, is the most significant gene affected by the ACHE/rs1799806 mutation.

**Table 4:** List of SNPs showing cis eQTL (p value < 0.001) from Brain and whole blood tissue using GTEx v8 database.

| CHR | SNP | Contribute to<br>Gene | p-value | Tissue |
| --- | --- | --- | --- | --- |
| 7 | rs1799806 | SLC12A9 | 3.00E-18 | Brain - Cerebellum |
| 7 | rs1799806 | SLC12A9 | 1.90E-15 | Brain - Cerebellar Hemisphere |
| 7 | rs1799806 | EPHB4 | 7.00E-14 | Whole Blood |
| 7 | rs1799806 | MOSPD3 | 1.70E-13 | Whole Blood |
| 7 | rs1799806 | UFSP1 | 3.30E-12 | Whole Blood |
| 7 | rs1799806 | GIGYF1 | 2.40E-09 | Whole Blood |
| 7 | rs1799806 | SLC12A9 | 3.50E-09 | Brain - Cortex |
| 7 | rs1799806 | SLC12A9 | 3.00E-08 | Brain - Caudate (basal ganglia) |
| 7 | rs1799806 | UFSP1 | 9.20E-07 | Brain - Nucleus accumbens (basal<br>ganglia) |
| 7 | rs1799806 | SLC12A9 | 0.0000011 | Brain - Nucleus accumbens (basal<br>ganglia) |
| 7 | rs1799806 | GIGYF1 | 0.0000023 | Brain - Nucleus accumbens (basal<br>ganglia) |
| 7 | rs1799806 | GIGYF1 | 0.000005 | Brain - Putamen (basal ganglia) |
| 7 | rs1799806 | STAG3 | 0.0000053 | Brain - Cortex |
| 7 | rs1799806 | SLC12A9 | 0.0000086 | Brain - Putamen (basal ganglia) |
| 7 | rs1799806 | PVRIG | 0.0000096 | Brain - Caudate (basal ganglia) |
| 7 | rs1799806 | STAG3 | 0.000013 | Brain - Cerebellum |
| 7 | rs1799806 | STAG3 | 0.000066 | Brain - Caudate (basal ganglia) |
| 8 | rs28686812 | ARC | 5.80E-11 | Whole Blood |
| 8 | rs2234911 | ARC | 2.80E-11 | Whole Blood |
| 4 | rs35172627 | DCHS2 | 0.000045 | Brain - Cerebellum |
| 6 | rs1570070 | RP11-288H12.3 | 0.000014 | Whole Blood |
| 11 | rs12453 | MS4A6A | 1.00E-18 | Whole Blood |
| 6 | rs367398 | HLA-S | 6.50E-08 | Whole Blood |
| 6 | rs367398 | HLA-DRB6 | 8.30E-07 | Whole Blood |
| 6 | rs367398 | XXbac-BPG181B23.7 | 0.000058 | Whole Blood |
| 6 | rs367398 | ZBTB12 | 0.00011 | Whole Blood |

ACHE gene located on chromosome 7q22.1 encodes for Acetylcholine esterase(AChE) enzyme was previously implicated as a risk factor for AD. Acetylcholine esterase commonly associated with Aβ plagues and NFT in AD[25]. Also, AChE inhibitors are widely used for the treatment of AD[26]. However, to the best of our knowledge, there is currently no literature on ACHE and its variants that have a role in Alzheimer’s disease. This is the first study showing the association with AD.

Also, apart from ACHE/rs1799806 variant NOTCH4/rs367398 variant is showing significant eQTL with multiple neighboring genes within Whole Blood with most significant eQTL for HLA-S gene with p value of 6.50E-08.

An intricate network of protein-protein interactions plays a vital role in the regulation of biological processes. A mutation in any of the genes in the network may also disrupt pathway[27]. Fig. 3 depicts a network analysis of significant genes. The ACHE gene has a tight link with the APP and PSEN1 genes. NOTCH4, on the other hand, has a significant link to APP and ADAM10. CLU has a strong link to the APOE, APP, and MAPT genes.

**Figure 3:**
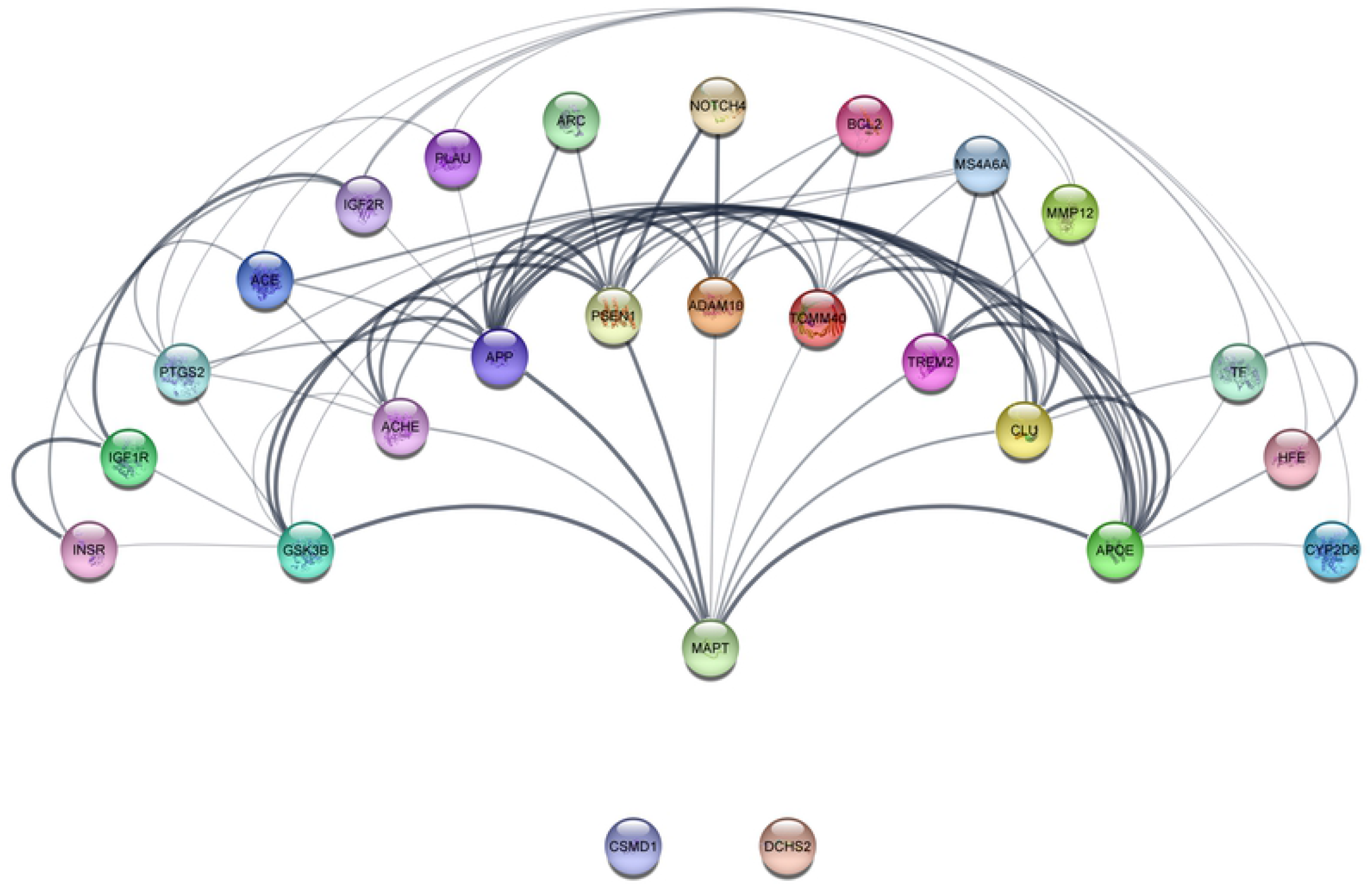
Gene-Gene interaction analysis of 16 significant genes in the present study with top 10 AD-associated genes selected from the Disgenet database based on gene disease association score.

Except for the DCHS2 and CSMD1 genes, all of the significant genes interact with AD-related genes either directly or indirectly linked to APP and MAPT gene functioning.

Studies have shown that the CSMD1 (CUB and Sushi Multiple Domains 1) gene plays a role in the regulation of complement activity and in inflammatory response, as well as in cognitive functions and synaptic plasticity[28]. Prolonged inflammatory responses have been reported in Alzheimer’s disease patients, and they are not only associated with neuronal loss, but they may also be responsible for aggravating Aβ deposition and neurofibrillary tangles[29]. DCHS2 gene’s (Dachsous cadherin-related 2 protein) exact function is still unclear, but it is associated with the progression of neurofibrillary tangles, which is directly correlated with cognitive impairment[30].

## 4. Conclusion

From this study we can conclude ACHE/rs1799806 and NOTCH4/rs367398 variant is showing most significant association with AD. However, no variants except NOTCH4/rs367398 has passed multiple test corrections. The main constraint of the present study is the small sample size, and none of the variants holds significance after employing multiple test corrections. Further validation with a large sample size is required to further implicate the role of these variants in AD pathogenesis.

## Acknowledgement

Authors wish to thank HMH-Ahmedabad and HMH-Bhuj for collection of blood samples & screening of Alzheimer’s patients. Authors are also thankful to DST (Department of Science and Technology) for providing necessary funding to carry out the study.

## Statement of Ethics

The Ethics Committee of HMH-Gujarat Institute of Mental Health approved the study.

## Author contributions

Conceived and designed the experiments: CJ AP. Performed the experiments: UB. Analysis and interpretation: UB AP CJ. Wrote the paper: UB. Sample collection and clinical screening of Alzheimer’s patients: AC DB CP VD.

## Data Availability

The datasets used in this article have been submitted to the National Center for Biotechnology Information’s Sequence Read Archive (SRA), where they can be accessible under the bioproject PRJNA824623 and accession number SRP368230.

## Supporting Information

S1 Table. Details of Regions Targeted

